# DeepDiveR – A software for deep learning estimation of palaeodiversity from fossil occurrences

**DOI:** 10.1101/2024.09.03.610960

**Authors:** Rebecca B. Cooper, Bethany J. Allen, Daniele Silvestro

## Abstract

1. The incompleteness of the fossil record, in particular variation in preservation and sampling through space and time, presents a barrier to estimating changes in biodiversity which standard statistical methods struggle to account for.
2. Here we present DeepDiveR, an R package for the DeepDive program enabling estimation of biodiversity from fossil occurrence data. The method uses a simulation-trained deep neural network to generate predictions of biodiversity change through time, while accounting for temporal, spatial and taxonomic heterogeneities in preservation.
3. DeepDiveR can be readily used to explore the extinct biodiversity of different clades. We demonstrate the pipeline to build and customise analyses, including consideration of changes in biogeography. We also further develop the model to integrate information about modern diversity in the case of extant clades and introduce a function that automatically adjusts the parameterization of the simulations to generate training data that reflect the distribution of empirical datasets.
4. To demonstrate the software, we analyse the fossil record of the order Carnivora through the Cenozoic, finding a peak in diversity in the Late Miocene and a 37% species loss since the Pleistocene. Our implementation includes the generation summary statistics and plots that allow for an evaluation of the model performance and diversity estimations and a configuration file that captures all parameters required to guarantee the full reproducibility of the results.

## 1 Introduction

Estimating how many taxa have existed in the past, how biodiversity has changed through time, and what mediates the diversity of life on Earth, has been a topic of major debate for many decades in palaeontology (Valentine *et al*., 1969; Raup, 1976; Valentine *et al*., 1978; Sepkoski, 1981; Benton & Emerson, 2007; Alroy, 2010; Ezard *et al*., 2011; Benton, 2015; Harmon & Harrison, 2015; Rabosky & Hurlbert, 2015; Benson *et al*., 2021), with much still to be resolved. Understanding diversity changes and their causes is essential to contextualising current and future changes in biodiversity, particularly as the fossil record contains the only direct evidence of response to drastic changes in Earth systems, such as climate change, at magnitudes which have not been observed on historical time scales (Jackson, 2010; Andermann *et al*., 2020; Dietl, 2019).

However, estimating biodiversity from fossil occurrence data remains a difficult task due to the inherent incompleteness of the fossil record. Less than 1% of species are thought to be preserved and then rediscovered by palaeontologists (Jablonski, 2004), with 30% of today’s tetrapods having virtually no potential to be recorded at all (Krone *et al*., 2024). Addressing the incompleteness of the fossil record is key to understanding how biodiversity has changed over life’s history. In recent years it has been demonstrated that while commonly used rarefaction methods (Alroy, 2010, 2020; Chao & Jost, 2012) can reliably correct for temporal variation in the quality of relatively well sampled fossil clades, they typically fail to account for the full range of heterogeneity in the fossil record particularly at the largest spatial scales (Close *et al*., 2020a,b). Growing appreciation of the impact that spatio-temporal biases have on the fossil record (Benson *et al*., 2021; Antell *et al*., 2024) has led to the development of new methods (Flannery-Sutherland *et al*., 2022b; Antell *et al*., 2024; Dunne *et al*., 2023; Thompson *et al*., 2020; Hauffe *et al*., 2022; Alroy, 2020) to correct bias in estimates, new databases of modern and palaeontological records that include a higher spatial grain have been compiled (Smith *et al*., 2023), greater understanding of the spatial variation which can result from choice of plate models (Buffan *et al*., 2023; Jones & Domeier, 2024) in geographic assignments, and development of spatial clustering protocols which are becoming common practice in untangling variation in organismal distributions through space (Daumantas & Spiridonov, 2024; Miele *et al*., 2014; Edler *et al*., 2016; Vilhena & Antonelli, 2015; Close *et al*., 2020a; Flannery-Sutherland *et al*., 2022b; Jones *et al*., 2023).

One recent method for estimating palaeobiodiversity from the fossil record using deep learning, named DeepDive (Deep learning Diversity Estimation), uses a coupled stochastic modelling and deep learning approach that accounts for variation in the temporal, spatial and taxonomic scope of the fossil record through simulationbased training, and is implemented in Python (Cooper *et al*., 2024). The DeepDive method has several advantages over other frameworks: data are not subsampled, allowing estimates to be made even for poorly recorded clades with maximal data usage; taxonomic biases are considered alongside spatial and temporal variation in the record; and estimates can be made accurately at the largest spatial scales rather than requiring separate estimates for subsampled regions as in alternative methods (Flannery-Sutherland *et al*., 2022b; Close *et al*., 2020a,b; Vilhena & Smith, 2013). DeepDive performs particularly well under conditions of pronounced spatial biases, and allows users to consider potentially widely-varying and complex interactions of biases in the fossil record while removing the need to explicitly parameterise analyses to encompass these.

The availability of deep learning libraries (such as TensorFlow (Abadi *et al*., 2015)) and computational efficiency of Python made it the most logical language in which to develop DeepDive. However, the R environment remains the preferred platform among palaeontologists, as shown by the extensive and growing development of R packages in the field (e.g. Varela *et al*., 2015; Reitan & Liow, 2019; Kocsis *et al*., 2019; do Rosario Petrucci *et al*., 2022; Jones *et al*., 2023), and this might inhibit the use of DeepDive by the wider scientific community. Here we present DeepDiveR, an R package which helps users to create input and configuration files ready for DeepDive analyses. These files can then be executed in a single line in either a terminal or command prompt window, without requiring the user to write any Python code. Aside from facilitating ease of use, this analytical pipeline also means that any assumptions made and all model settings are recorded in a single file thus helping users to build customisable and reproducible workflows. The DeepDiveR package enables researchers interested in the diversity dynamics of a range of clades, particularly those concerned about fossil record bias, to investigate biodiversity patterns using a deep learning approach. We demonstrate use of the software in an analysis of the mammalian clade Carnivora.

## 2 Description

The DeepDiveR package automatizes the data preparation and the implementation of an analytical pipeline in DeepDive, which typically implies three steps: 1) simulation of training datasets, 2) training of deep learning model, 3) prediction of empirical diversity trajectories (Fig. 1). The DeepDive programme generates simulated biodiversity and biogeographic histories through space and time, which are then degraded to emulate the fossil record. These simulated data are then used to train a recurrent neural network model based on long short-term memory (LSTM) that learns to reconstruct the original biodiversity curves from the degraded fossil simulations. The LSTM is then used to estimate biodiversity through time from the known fossil record, even in the context of strong spatial biases. The DeepDiveR package generates pre-formatted input data and a configuration file containing all of the necessary settings to execute a DeepDive analysis.

**FIGURE 1:**
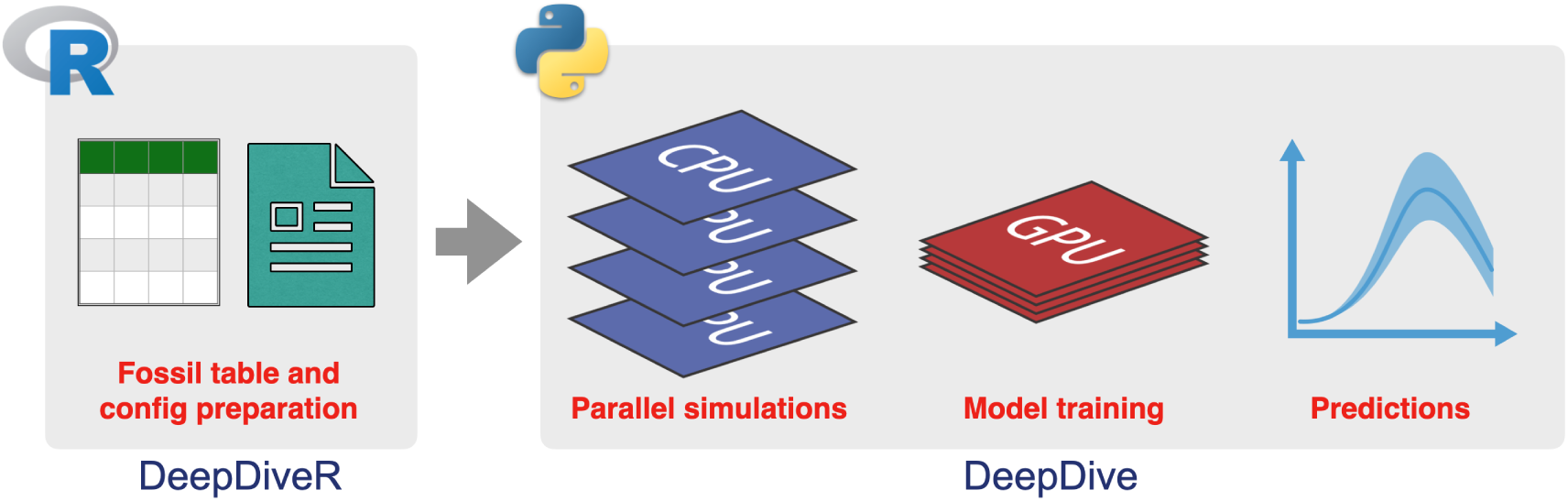
The DeepDiveR workflow. Firstly, in R, input data are preprocessed and prepared in the correct format for reading into DeepDive along with generation and editing of a configuration file (.ini) that stores all settings of the analysis. The config file is then executed in a command line terminal and runs in Python.

Steps can be taken to speed up the completion time of a DeepDive analysis. The simulation module can be efficiently parallelised on multiple CPUs, with computing time decreasing roughly linearly with the number of allocated CPUs (i.e. 100 CPUs will complete a given set of simulations *≈*100 times faster than if only one CPU is allocated). Models can be trained on a GPU for the most efficient computing times, if available, or will default to multiple CPUs otherwise.

The main outputs of the analytical pipeline include the trained models, the simulations, and a plot of diversity through time. Summary statistics describing the performance of the trained model (e.g. mean squared error on a test set) are saved in a .csv file. The output also includes a .csv file with estimates of biodiversity through time across age assignment replicates, that can be easily read in R for further plotting and analysis.

### 2.1 Installation and implementation

The development version of DeepDiveR can be run in R (v. 4.3.2 or above) and installed from a GitHub repository using the remotes R package (Csárdi *et al*., 2024):

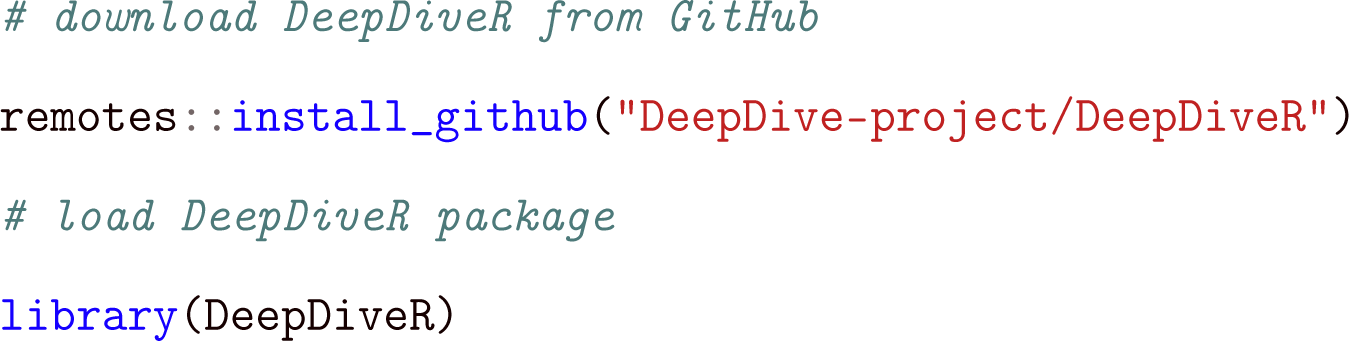

Once the input files have been created, DeepDive, which requires Python v.3.10 or above, can be installed using: python -m pip install git+https://github.com/DeepDive-project/deepdive

(in Windows use py instead of python). For further details on installation see https://github.com/DeepDive-project. The script run_dd_config.py available on GitHub is used to launch analyses.

### 2.2 Functions

DeepDive requires two files in order to run an analysis: the input file, which includes the fossil occurrence data on which the analysis is run, and the configuration file, which includes all the settings used during the analysis. There are two key functions in DeepDiveR that prepare these files, as explained below.

#### 2.2.1 Data input file

The function prep_dd_input takes a data frame of fossil occurrences (see Table 1) that contains one row per occurrence with columns specifying the taxon name, spatial information (e.g. continent or ocean basin and locality ID) and the minimum and maximum ages of its stratigraphic range. The output of the function is a .csv file containing the boundaries of user-defined time bins that will be used in the analysis, the number of localities sampled per discrete sampling region per time bin, and counts of occurrences for each taxon in each time bin per region. Fossil occurrences are assigned a point age, which by default are sampled from the respective stratigraphic age range of each locality The age sampling can be repeated a number of times (set through the r argument) to account for age uncertainties that will be reflected in the resulting diversity trajectories.

**TABLE 1:** An example input dataset, here of Cenozoic carnivorans, after cleaning. In absence of locality names or identifiers these can be allocated, as is done here, by assigning a number to each occurrence with the same MinAge, MaxAge, Latitude and Longitude. The “Area” column should indicate the discrete regions intended for use in the DeepDive analysis, in this case continents.

| Taxon | Area | MinAge | MaxAge | Locality |
| --- | --- | --- | --- | --- |
| Ailurus | Asia | 0.6 | 1.3 | 1 |
| Alopecocyon | Europe | 5.3 | 7.1 | 2 |
| Eucyon debonisi | Europe | 5.3 | 7.1 | 2 |
| Vulpes stenognathus | North America | 7.7 | 7.8 | 55 |
| Dinocrocuta algeriensis | Africa | 5.3 | 7.1 | 3289 |
| Acinonyx jubatus | Africa | 3.5 | 3.8 | 880 |
| Smilodon gracilis | North America | 1.9 | 2.5 | 700 |
| Smilodon gracilis | South America | 0.5 | 1 | 1879 |
| Arctotherium wingei | South America | 0.012 | 0.126 | 3610 |
| ⋮ | ... | ... | ... | ... |

Fossil occurrence data can be readily obtained from many large online databases such as the Paleobiology Database (Uhen *et al*., 2023), the New and Old Worlds (NOW) database (Žliobaitė *et al*., 2023), NEPTUNE sandbox (Renaudie *et al*., 2020), BioDeepTime (Smith *et al*., 2023) and Neotoma (Williams *et al*., 2018). Scrutiny is essential when these data are used, however: databases may contain unresolved or conflicting taxonomic assignments, use different palaeogeographic reconstructions to assign coordinates to occurrences, and most are missing substantial quantities of data, for example many known specimens remain unpublished in museums (Marshall *et al*., 2018). Prior to input file creation, it is essential that the fossil occurrence dataset is thoroughly checked and cleaned. Many computational tools exist to aid with data cleaning, including R packages such as CoordinateCleaner (Zizka *et al*., 2019), fossilbrush (Flannery-Sutherland *et al*., 2022a) and palaeoverse (Jones *et al*., 2023). Input data which may need to be cleaned include taxonomic identifications, age estimates of fossil occurrences and geographic assignments.

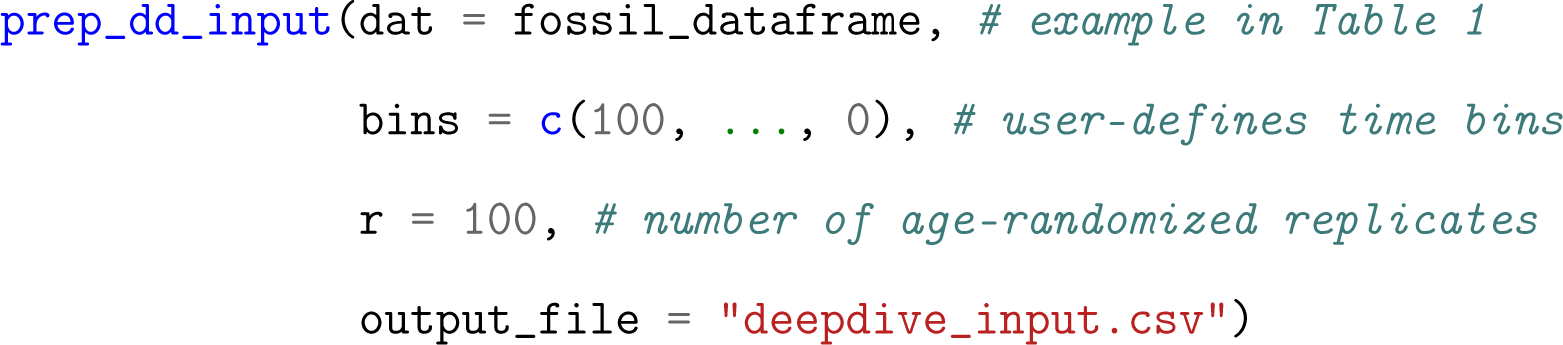

#### 2.2.2 Configuration file

The function create_config generates a text file (with .ini extension) using the ConfigParser package in R (Hoefling, 2017). The file includes a range of settings used in the DeepDive analysis, in a format which is both human-readable and editable.

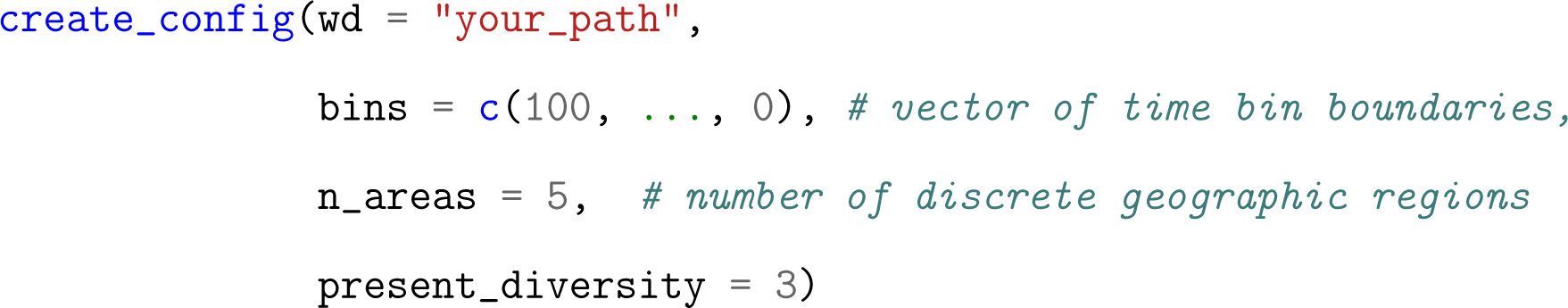

The configuration file contains default values for most parameters, and is split into four blocks. The first, general, contains settings that will be referenced throughout the workflow, such as file paths, data, time bins and number of discrete sampling regions, which will always need to be provided. The next blocks follow the steps of the method, namely simulations contains settings for defining the parameter space in training simulations of biodiversity with biogeographic ranges and the degradation of these data in the fossilisation and sampling simulator, model_training defines the parameters related to training the LSTM model to estimate biodiversity through time from the degraded data, and empirical_predictions provides settings for estimating biodiversity from empirical fossil occurrence records.

These modules can be run individually or in any combination given the required inputs are provided, so that workflows can be saved, evaluated and rerun when making adjustments to parameterisation. Once the config object has been created, the parameter values described can be manually adjusted using the set_value function (for instance the example below sets the number of training simulations to 10000):

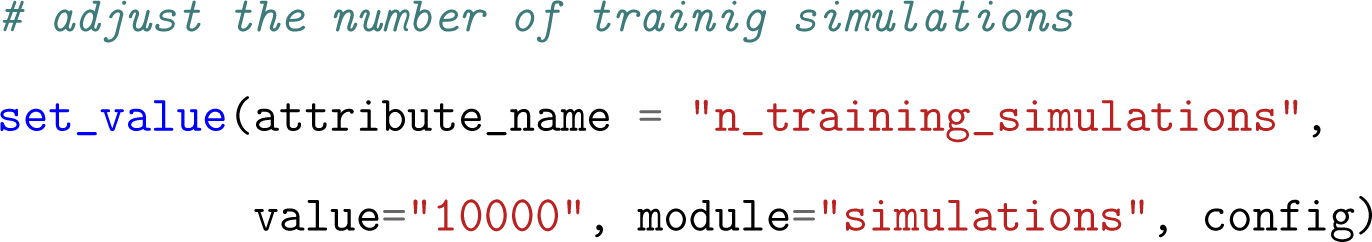

To simulate scenarios where regions only become available to a clade through time, the function areas_matrix can be used where area_ages is a matrix of minimum and maximum age estimates for establishing a connection between regions. If label is set to “end” regions will disappear in the specified age range in the simulations. In the following example, the first four areas are available from the start of the time window of interest (here the past 66 myr) while the last area only becomes available sometime between 12 and 7 Ma, for instance reflecting the emergence of a new land bridge between continents.

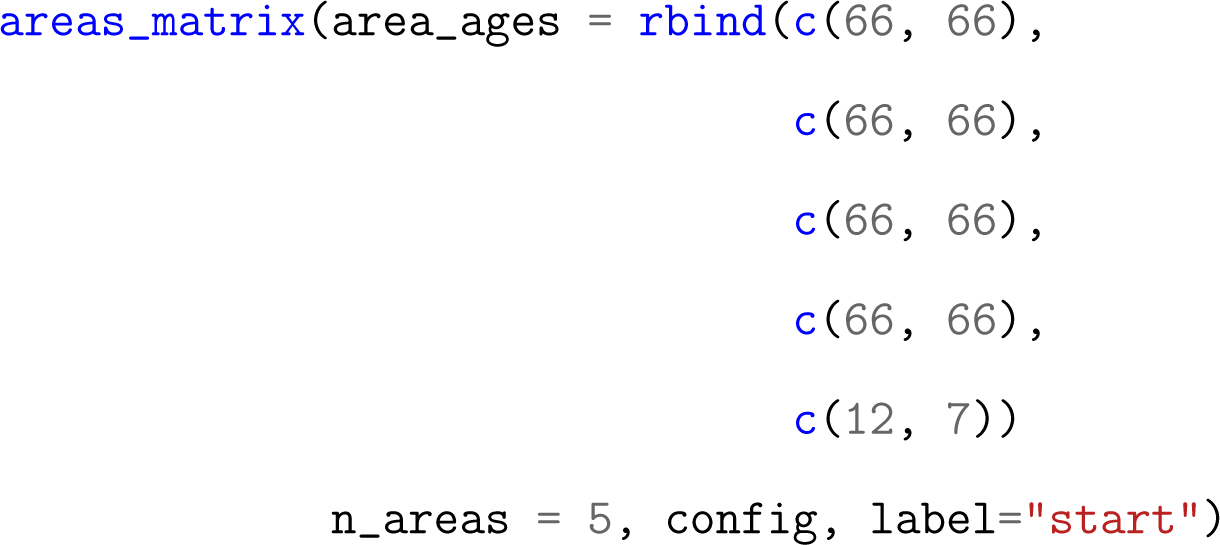

The configuration file can then be saved:

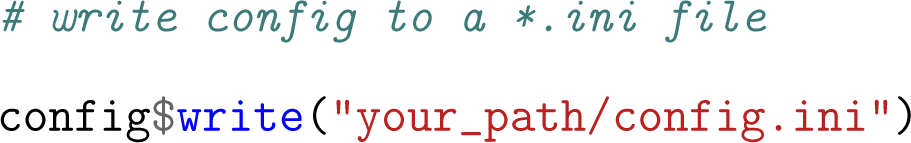

#### 2.2.3 Running a DeepDive analysis

Once the configuration and input files are created, the full DeepDive analysis (Figure 1), inclusive of simulation, model training and empirical predictions, can be carried out through a simple command line entered in a Terminal (MacOS and Linux) or Command prompt (Windows) window using the Python script run_dd_config.py:

python run_dd_config.py your_path/config_file.ini

The user can additionally specify a working directory where all output files are saved and the number of CPUs used for the parallelized simulations, which will overwrite the corresponding settings in the configuration file, using the flags -wd your_working_directory and e.g. -cpu 64.

This command will create a “simulations” folder containing the training and test sets, and a “trained_models” folder containing the trained models and plots of the training history. The latter will additionally include plots comparing the empirical and simulated fossil features (e.g. number of occurrences through time and per region, number of localities, fraction of singletons, and sampled diversity), CSV files with the predicted diversity trajectories for the test set and for the empirical dataset, and a plot of the estimated diversity trajectory (Figure S1).

### 2.3 DeepDive model developments

Alongside DeepDiveR, here we also present two new features which build on the DeepDive implementation of Cooper *et al*. (2024): 1) the ability to autotune the simulation parameters to better reflect an empirical dataset, and 2) the ability to condition the model based on extant diversity.

#### 2.3.1 Autotuning simulation parameters

As DeepDive models are trained on simulated datasets it is important that these cover a wide range of evolutionary and preservation scenarios within which the empirical data is likely to be found (Cooper *et al*., 2024). A way to evaluate this is by looking at the range of simulated features (e.g. number of occurrences per area and through time, number of localities, etc.) and ensuring that the corresponding empirical features fall within that range. Previously, manual manipulation of the configuration settings via trial and error was required to ensure that this was the case. However, these parameters can now be automatically adjusted when running the analysis in DeepDive through an autotune function implemented in the Python library. This function runs in the background when executing the configuration file and changes the parameter values so that the resulting simulated datasets are similar to the empirical dataset. This will, for instance, affect the number of simulated sampled species, their average longevity, the frequency of singletons, the presence of gaps in the data, and differences in the number of records sampled across regions. The adjusted parameter values are automatically saved in a new configuration file to ensure full reproducibility of the simulations and subsequent analyses and to allow for manual inspection of the selected values. We recommend using autotune for empirical analyses as it ensures the model is appropriately parameterised, reducing the sources of error and the amount of guesswork required by users.

### 2.4 Conditioning on modern diversity

We also extend the DeepDive model to account for modern diversity, i.e. the number of living taxa in a clade. To use this feature, in the general block of the configuration file the variable present_diversity is set to the number of known extant taxa. If present_diversity is provided (replacing the default NA value) then predictions of diversity trajectories are inferred while considering this value as known. Specifically, this means that modern diversity will be included along with the fossil features in the input feeding into the LSTM. As a consequence, during training, the model learns that this feature contains important information about diversity at the end of the time interval. As we use bidirectional LSTMs, the output of the long short-term memory units projects this back through time and estimates of diversity leading up to the present will also be adjusted thereby conditioning time bins before the present. Once the LSTM is trained and the empirical analysis stage of the workflow begins, the estimate of present_diversity provided in the function create_config will be entered as this additional feature to condition empirical estimates.

In simulated test sets we find that conditioning estimates on modern diversity reduces errors by almost 40% (mean squared error MSE unconstrained = 0.388, MSE conditioned = 0.244) in the case of simulations of extant clades (Figure 2, Figure S2). Interestingly, even in extinct clades, i.e. when present_diversity = 0, this conditioning leads to some accuracy improvement, albeit of a smaller magnitude of *≈* 10% (MSE unconstrained = 0.363, MSE conditioned = 0.328). Conditioning on modern diversity reduces errors in time bins prior to the present, although the impact of this conditioning appears to have longer duration when the simulated clade is extant (Figure 2C-D). The impact on errors does not appear to be uniform across simulations, although a general improvement is seen (Figure S3).

**FIGURE 2:**
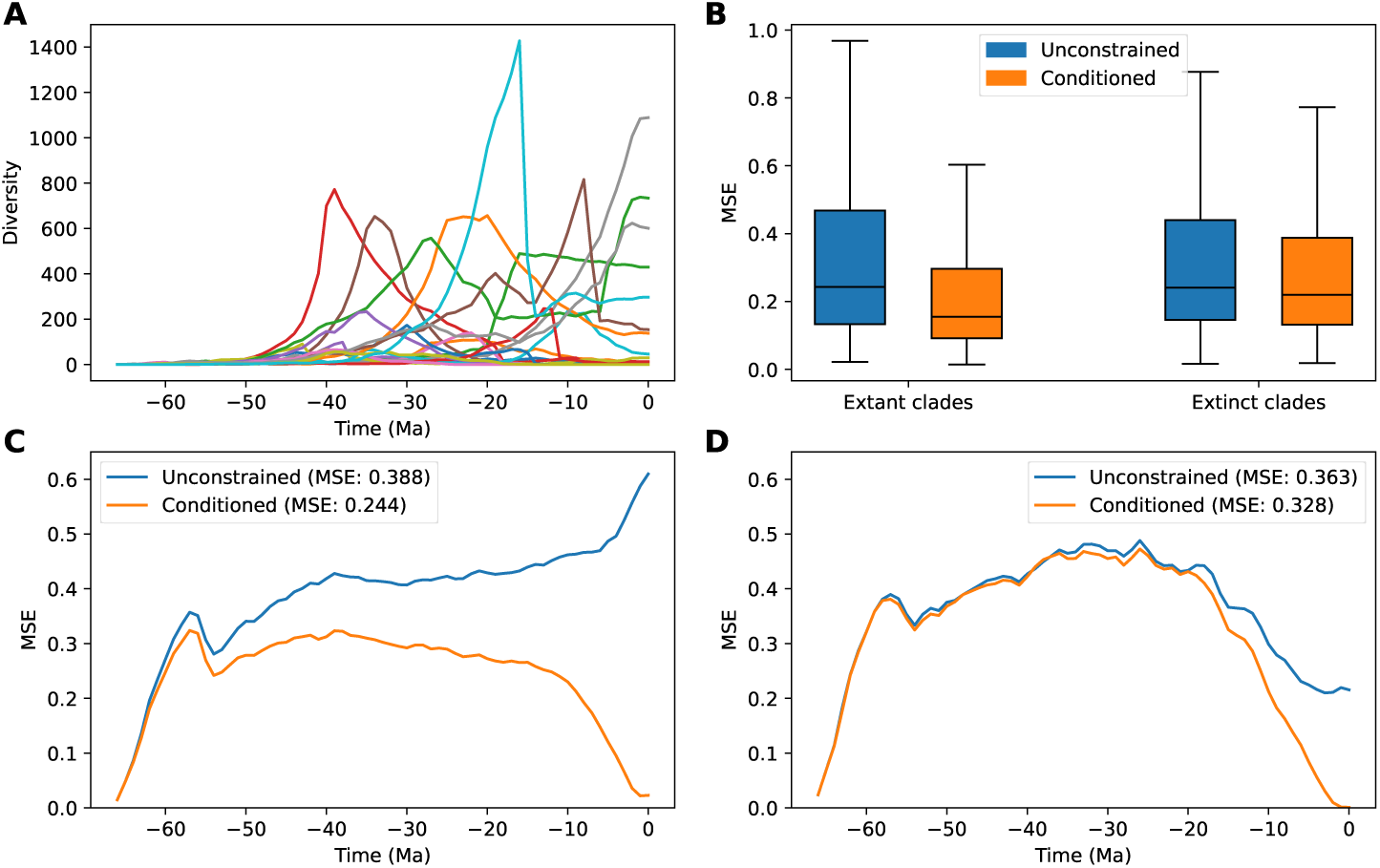
Performance of model on test sets with and without conditioning on modern diversity. A) Examples of simulated diversity trajectories in a training set. D) MSE for a model trained without constraints on modern diversity (blue) and for a model trained while conditioning on modern diversity (orange) for test sets of extinct and extant clades. C) MSE through time for the mean of simulated test sets where the clade remains extant at the end of the time frame when the model is unconstrained (blue) or conditioned on modern diversity (orange), N=3310. D) MSE through time for the mean of test simulations where the clade is extinct by the end of the time frame when the model is unconstrained (blue) or conditioned on extant diversity i.e. 0 (orange), N=1690.

## 3 Example application: Cenozoic carnivorans

To demonstrate the DeepDiveR analytical pipeline, we analyse the global record of Cenozoic mammalian carnivores including stem carnivorans (Faurby *et al*., 2019), comprising 13,006 occurrences across 1,574 species. We generated a configuration file to simulate a training set with clade origination between 66 and 52.8 Ma, and where one of the areas (South America) could only be colonized starting from 11.608 and 7.3 Ma, reflecting the known dynamics of the Great American Biotic Interchange (Bacon *et al*., 2015).

Localities were assigned to groups of unique occurrences sharing the same age range and palaeocoordinates rounded to the nearest 0.1 km. Autotune was used to adjust the settings used for simulations, to ensure that the range of the empirical data fell within the scope of simulations the model is trained to work with, and we conditioned the analyses on a modern diversity of 313 living species (Faurby *et al*., 2019). We randomly resampled fossil ages 100 times and trained four models with different architectures (Figure S4): 2 LSTM layers of 64 and 32 nodes; 3 LSTM layers of 128, 64 and 32 nodes; 3 LSTM layers of 256, 128 and 64 nodes; 2 LSTM layers of 512 and 128 nodes. The randomization and the different models allowed us to infer carnivore diversity through time along with a confidence interval.

Our analysis found that carnivoran diversity climbed steadily in the fist 15 My from the origin of the clade (Figure 3), reaching around 350 species towards the end of the Ypresian, during the Early Eocene Climatic Optimum. Diversity then dropped by around 60% by the Late Eocene around 40 Ma. This decline in diversity may be related to a shift to cooler climates (Bohaty & Zachos, 2003). After the Eocene-Oligocene boundary diversity began to accumulate again more steeply, reaching around 400 species by the end of the Oligocene. After a further interval of more modest diversity accumulation in the early Miocene, the diversity remained stable with some fluctuation for almost 20 myr, leading to an estimated *≈*500 species at the beginning of the Pleistocene. Carnivore diversity entered a phase of rapid decline in the late Pleistocene, with species numbers decreasing to the 313 extant species, representing recent loss of approximately 37% of biodiversity. While the absolute estimates come with a large uncertainty interval, the trends hold consistent across model architectures and age randomisations.

**FIGURE 3:**
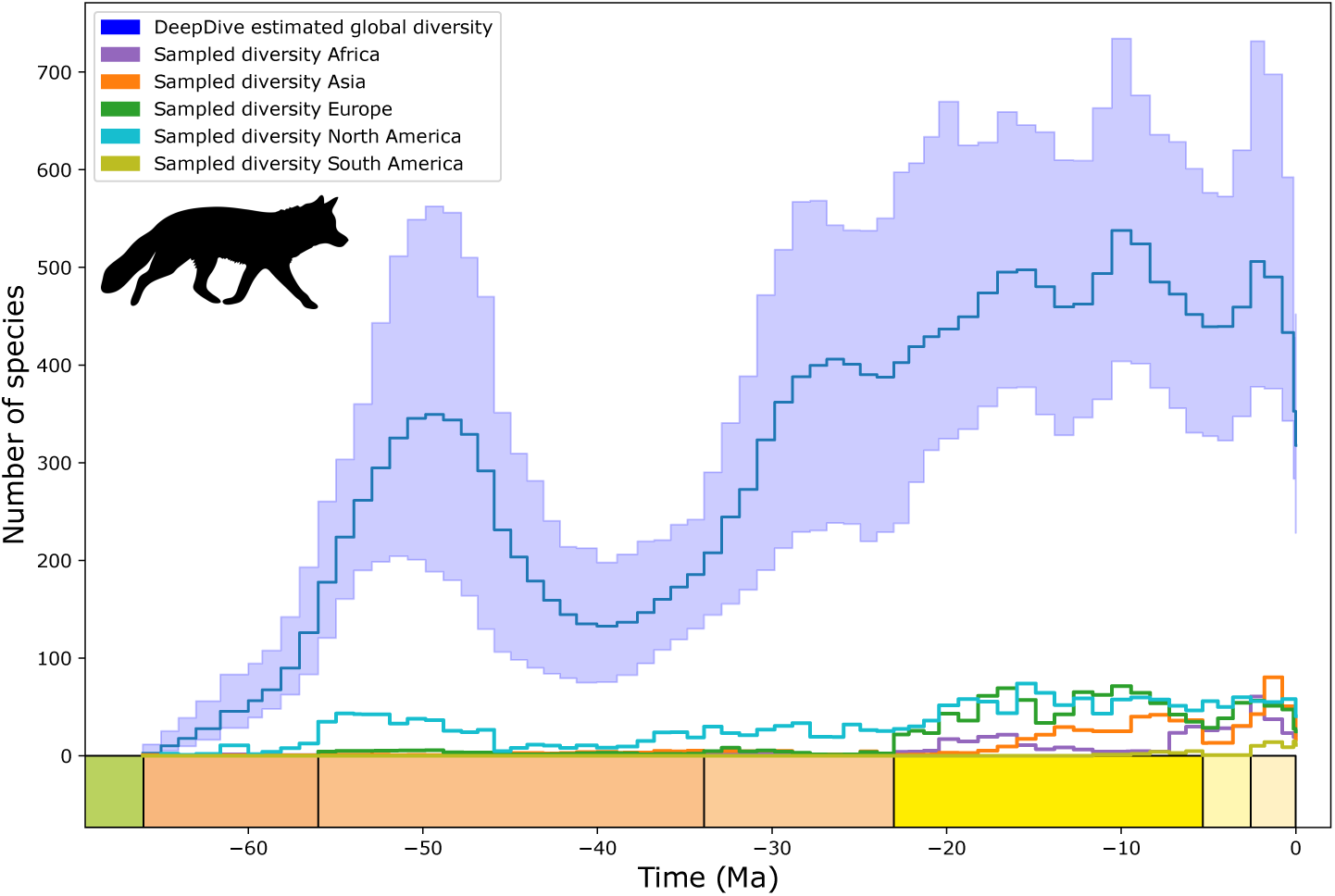
Global carnivoran diversity at the species level for the last 66 My estimated using DeepDiveR (blue). Shaded blue intervals indicate estimates made across an ensemble of four model architectures with similar performance (MSE = 0.259–0.262). Most of the study interval sees accumulation of genera, with losses in net diversity occurring during the shift to cooler climates after the Early Eocene Climatic Optimum and from the Pliocene onwards, after a Miocene peak in diversity. Diversity is stable during the latter halves of the Eocene, Oligocene and for most of the Miocene. Sampled diversity for each continent: Asia (orange), Europe (green), North America (light blue), Africa (purple) and South America (yellow).

The trends we observe here are broadly consistent with recent phylogenetic estimates of the diversity of carnivorans through time and space (Faurby *et al*., 2019), which however do not fully account for unsampled species. This might explain that fact that our estimates exceed phylogenetic ones. We also find some differences in diversity trends, for example that diversity accumulates towards its peak earlier than in phylogenetic estimates, beginning around 30 Ma, before remaining somewhat stable for longer. This increase in diversity around 30 Ma has also been observed in studies of the North American fossil record as replacement communities with more carnivoran species rise following losses of previous faunas during the cooling phases of the Middle and Late Eocene (Van Valkenburgh, 1999; Janis, 1993). Indeed the underlying diversity of dental adaptations for feeding in mammalian carnivores still seen today has been established by this time with dental diversity modulated by the availability of prey (Van Valkenburgh, 1988), a relationship that is often invoked to explain the relationship between climate changes and the indirect response of carnivoran diversity via limited food resources throughout the history of the clade and today (Sandom *et al*., 2013; Nascimento *et al*., 2024; Wolf & Ripple, 2016). The general stability of carnivoran diversity throughout the past 20 myr with upticks in diversity after the Mid-Miocene climatic Optimum and on entering the Pliocene is consistent with estimates of net diversification rates inferred in previous analyses of the fossil record (Liow & Finarelli, 2014). However, these trends are inconsistent among continents (Finarelli & Liow, 2016; Pires *et al*., 2015; Hauffe *et al*., 2022) and although carnivore guilds are potentially near equilibrium at the global scale at this time, this is also a period of much turnover. Fluctuations in carnivoran diversity in the past 20 myr may be related to community restructuring possibly following the loss of prey species in the Miocene and Pliocene, as has been observed in studies within Europe where trophic restructuring generated a higher predator to prey ratio with gradual increases in carnivoran diversity despite transitions in faunas (Nascimento *et al*., 2024; Blanco *et al*., 2021). Additionally we observe a much sharper drop towards present day diversity at the end of the study interval. Recent substantial extinctions of mammalian carnivores are likely the result of a negative impact of modern humans and possibly earlier hominins through direct hunting competition for prey species, and habitat degradation (Ripple & Van Valkenburgh, 2010; Werdelin & Lewis, 2013; Faurby *et al*., 2020). This recent human-driven diversity drop is also consistent with strong biodiversity losses across several other mammalian clades (Andermann *et al*., 2020), and in line the the currently elevated extinction risk affecting many living species of carnivores (Red List, 2024).

## 4 Conclusion

The DeepDiveR software provides access to a simulation based deep learning programme previously only available to Python users, while making key functionalities simpler to utilise. The extended model includes conditioning on modern diversity, which can substantially increase the accuracy of the predicted diversity trajectories. We also introduce an autotune option to ensure the parameterisation of the simulations properly reflects the distribution of empirical data, and demonstrate these new features in an analysis of the clade Carnivora. These steps to improve the performance and accessibility of DeepDive pave the way for analyses of diversity in the fossil record of more clades, which is essential to addressing the knowledge gap that has been generated by variation in the completeness of the fossil record.

## Acknowledgements

We thank Sara Varela and Tiago B. Quental for their feedback and comments on the DeepDiveR project. RBC and DS received funding from the Swiss National Science Foundation (PCEFP3_187012). DS also received funding from the Swedish Research Council (VR: 2019-04739). BJA received funding from ETH Zurich.

## 5 Data availability statement

All scripts and data are open-source available on GitHub https://github.com/DeepDive-project. The DeepDiveR package is open-source and available at https://github.com/DeepDive-project/DeepDiveR, the DeepDive package is also open-source and available at https://github.com/DeepDive-project/deepdive. Carnivoran data scripts are available at https://github.com/DeepDive-project/example_files/tree/main/carnivoran_analysis.

## 6 Supplementary Figures

**FIGURE S1:**
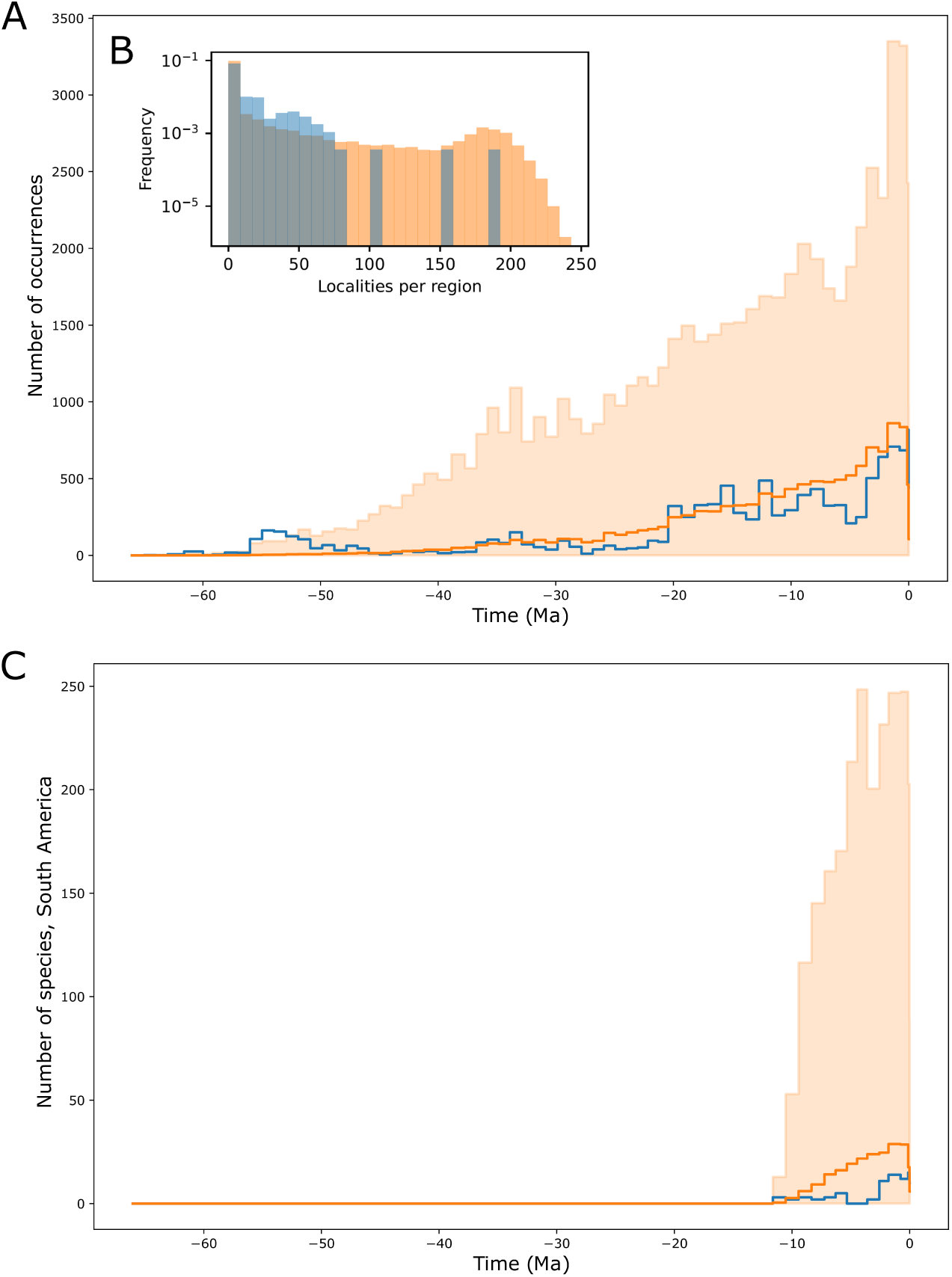
Examples of simulated features (orange) and empirical features (blue) from carnivoran analysis where A) shows the number of occurrences through time and B) is the frequency of localities per sampling region. C) The number of species through time in South America is shown as an example where areas are added through time using the areas_matrix function, in this case reflecting the closure of the Isthmus of Panama (Bacon *et al*., 2015).

**FIGURE S2:**
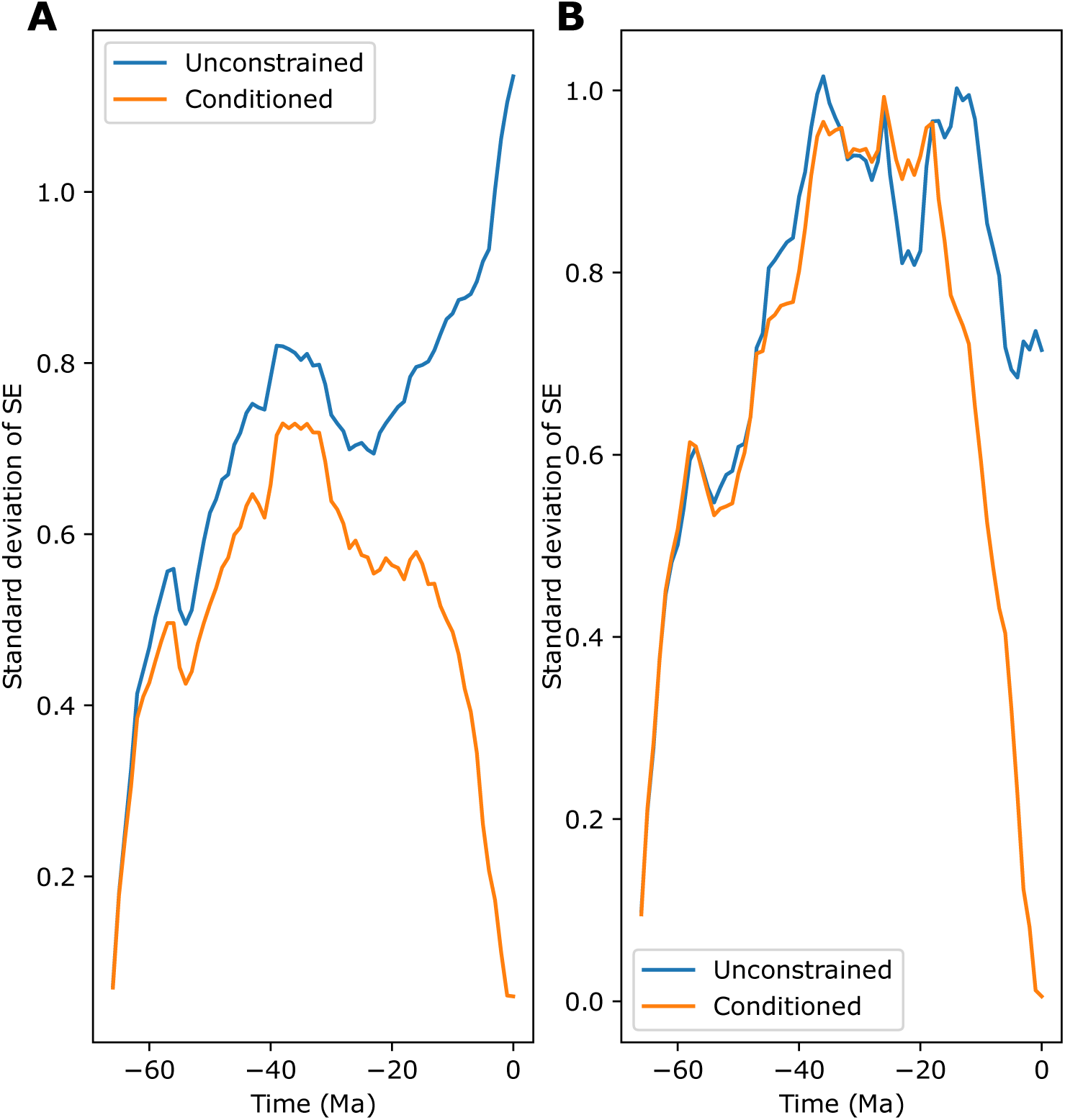
Standard deviation of mean squared errors through time (Ma) where models are unconstrained (blue) or conditioned on modern diversity (orange) for A) test simulations of extant clades (N=3310) and B) test simulations of extinct clades (N=1690).

**FIGURE S3:**
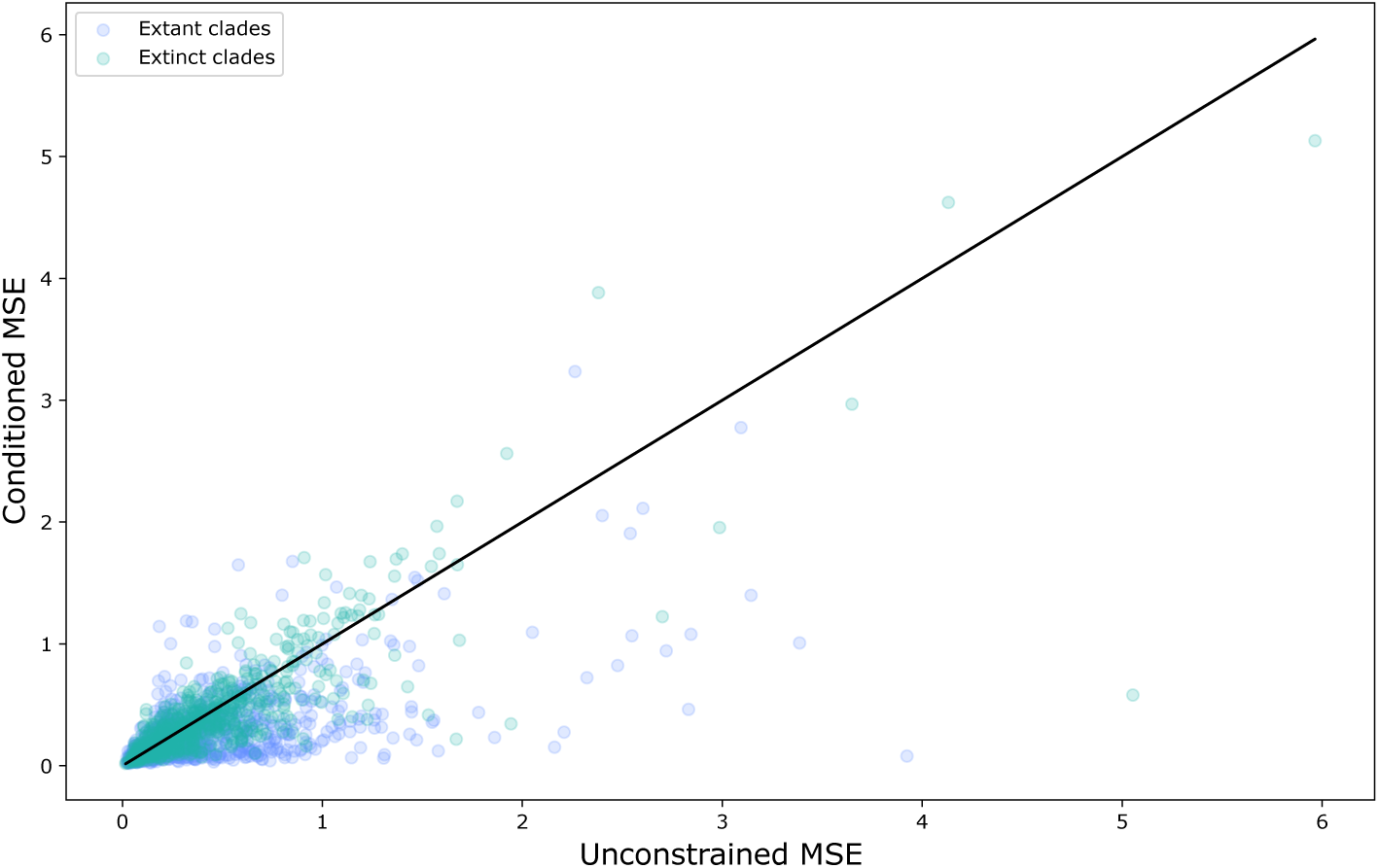
Mean squared error for test simulations where estimates are conditioned on modern diversity relative to MSE obtained where the model is unconstrained for the same test simulations. There is no visible trend in the effect of conditioning when the clade is extinct (green) or extant (purple).

**FIGURE S4:**
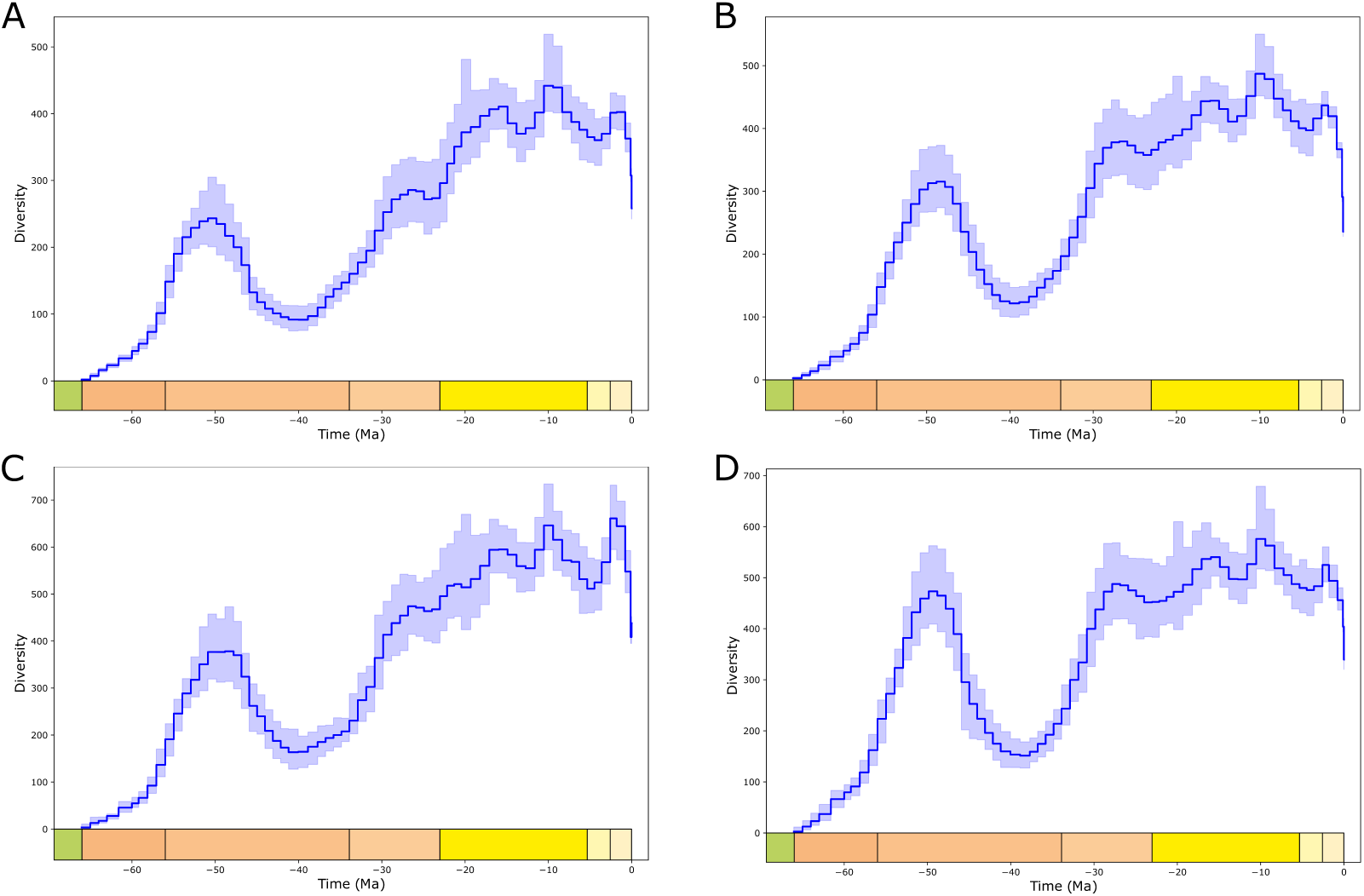
Global carnivoran diversity at the species level for the last 66 myr estimated using four model architectures in DeepDiveR with similar performance. All models contain 64, 32 dense node layers and between two and three LSTM layers with different number of nodes: A) 64, 32; B) 128, 64, 32; C) 256, 128, 64 and D) 512, 128.

## Notes

https://github.com/DeepDive-project/DeepDiveR

https://github.com/DeepDive-project/example_files/tree/main/carnivoran_analysis

### Competing Interest Statement

The authors have declared no competing interest.

